# Adaptive sampling in nanopore sequencing for PCR-free random access in DNA data storage

**DOI:** 10.1101/2024.11.05.622081

**Authors:** Roman Sokolovskii, Robert Ramirez-Garcia, Thomas Heinis

## Abstract

Adaptive sampling is a unique feature of nanopore sequencing—it enables the user to selectively enrich the sample (i.e., increase the effectively observed concentration of a subset of DNA in the sample) based on a user-provided set of reference sequences to target (or to avoid). It also enables dynamic switching of random access targets during a sequencing run as well as re-archiving of the sampled aliquot for later sequencing. As we demonstrate experimentally, the advantages of adaptive sampling make it a promising component of random access solutions in DNA data storage. We demonstrate that this approach allows selective data retrieval from a DNA pool without PCR, enabling the remaining aliquot to be re-archived for future retrieval of different target files. Our method helps to limit pool depletion effectively. It also allows dynamic switching between random access targets/data during a sequencing run. Adaptive sampling, however, primarily works for long sequences while for DNA data storage short oligos are typically used. We also show how to work around this limitation.

## Introduction

Most proof-of-concept DNA data storage systems currently store files on short DNA oligos (where one file is stored across multiple oligos) and use PCR amplification to provide selective (random) access to individual files stored in the DNA pool. Oligos belonging to the same file are equipped with the same PCR primers and random access to an individual file is implemented using a PCR amplification based on these primers [1]. PCR-based random access, however, has two major drawbacks: pool depletion and PCR bias.

Pool depletion means that every read operation requires an aliquot of the DNA pool. The aliquot will contain DNA that encodes information from all files in the pool, not just those requested by the user. The DNA that corresponds to the requested `file`(s) will be PCR-amplified, whereas the rest will be diluted away, and the information stored in those oligos will be irrecoverably lost. To minimise pool depletion, the system designer must either increase the number of physical copies of each oligo in the pool, increasing the cost of DNA synthesis and compromising the density advantage of DNA data storage, or decrease the aliquot size, exacerbating the second drawback of PCR-based random access—PCR bias.

PCR bias refers to the uneven representation of targeted oligos in a PCR-amplified sample. Both the inherent randomness of the amplification process and systematic dependencies of polymerisation rates on the contents of the amplified DNA mean that different oligos will be amplified at different rates, which, given the exponential nature of PCR amplification, may in extreme cases result in some oligos dominating the amplified sample while others are diluted away, hindering information recovery [2]. PCR bias becomes more acute as the starting pre-PCR oligo concentrations become more uneven or the number of PCR cycles increases, and both are likely as the aliquot size is decreased.

This work aims to circumvent the limitations of PCR-based random access by leveraging the strengths and unique features of nanopore sequencing technology, namely its ability to perform PCR-free sequencing, its adaptive sampling feature for random access, and the existence of library recovery protocols to reduce pool depletion.

Unlike PacBio [3] or Illumina [4] sequencing platforms, which include repeated DNA polymerisation as part of their sequencing process, nanopore sequencing does not inherently require PCR amplification during sample preparation or sequencing. This means that PCR bias becomes solely a consequence of using PCR amplification for random access, and using other random access mechanisms could eliminate the problem of PCR bias entirely and mitigate pool depletion.

Nanopore sequencing devices produced by Oxford Nanopore Technologies (ONT) potentially provides just such a mechanism, namely adaptive sampling. During adaptive sampling, the sequencing device performs real-time basecalling and aligns the readouts against a provided reference. If the match is observed, the device will continue sequencing the strand as it passes through the pore, otherwise sequencing stops and the strand is ejected i.e., returned to the aliquot, see Figure 2. To the best of our knowledge, adaptive sampling has not been investigated as a random access mechanism in DNA data storage applications.

**Fig 1.**
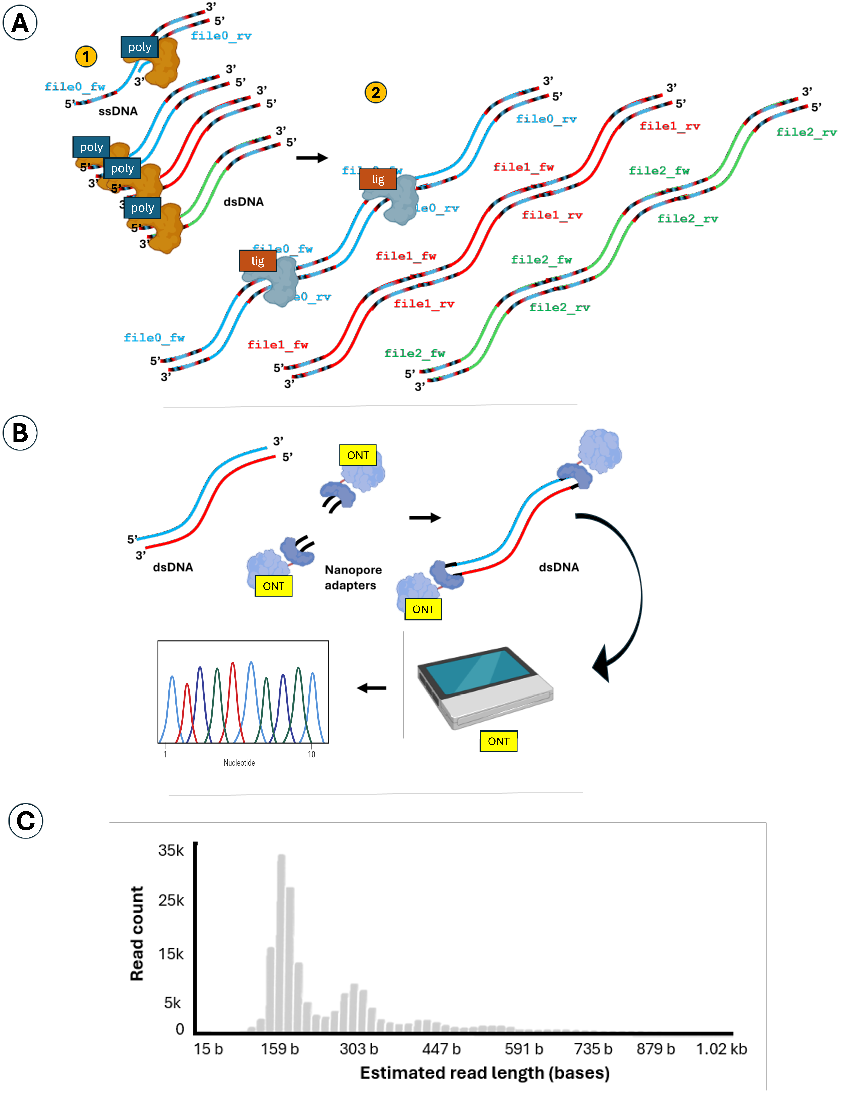
Sample preparation for ONT sequencing. **A)** DNA molecules belonging to the same `file` are elongated using a DNA polymerase and a DNA ligase (1)the 150-mer ssDNA pool is PCR amplified with specific primers corresponding to `file0`, `file1` or `file2`, creating three dsDNA pools where the files are separated; (2) a DNA ligase is employed to `file`-specifically conjoint the 150-mer dsDNA fragments into longer ones. **B)** The elongated double-stranded DNA molecules are adjoint to ONT-specific adapters for ONT sequencing before being mixed in proportions before ONT sequencing. **C)** Estimated read length count after ligation as measured during ONT sequencing.

**Fig 2.**
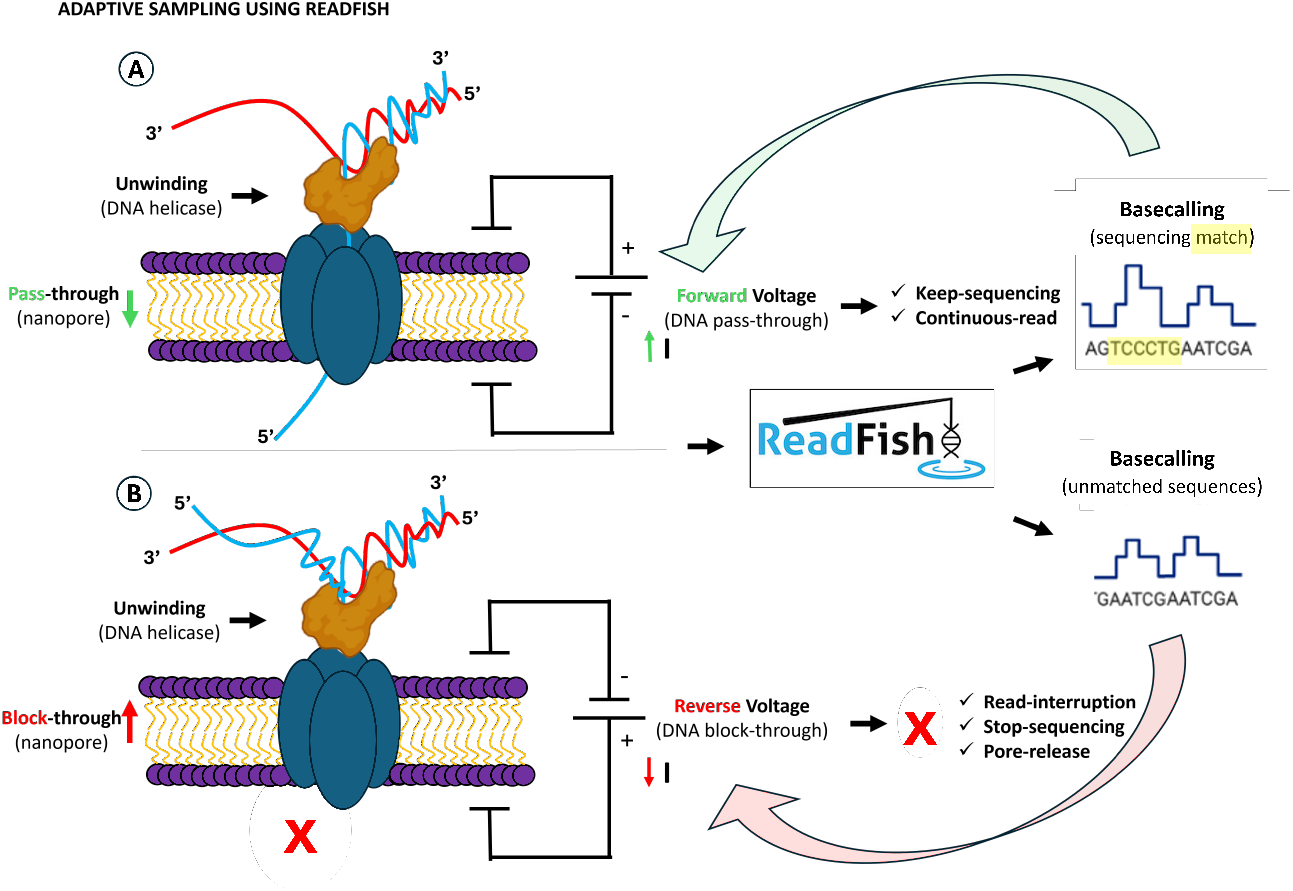
Adaptive sampling forward-or-reverse voltage switching with nanopore technology. **A)** Upon sequencing matches. **B)** After unmatched sequencing.

In addition to avoiding PCR bias, adaptive sampling for random access enables re-using some of the sequenced sample by employing a library recovery protocol (Methods, M5). The recovered library will contain parts of the pool that were not requested by the user. These will be available for subsequent sequencing if requested at a later stage, limiting pool depletion. Notably, re-using the library in this fashion is not possible after PCR amplification because in this case the non-requested oligos are diluted away.

The major obstacle to the use of adaptive sampling is that processing delays in real-time basecalling and alignment mean that by the time a decision on whether to keep sequencing a strand is made, about 800 nucleotides will already have passed the pore (400 nucleotides worth of ionic current data are buffered before the start of basecalling [5], with further delays in subsequent processing steps), whereas current DNA synthesis is limited to generating oligos of about 200-300 nucleotides in length (longer gene-length products can be assembled from short oligos; we expect that the cost and effectiveness of such methods will be brought down over time [6]). This means that, prior to sequencing, the oligos belonging to the same file will need to be assembled into longer strands, using standardized DNA ligation protocols. An additional advantage of sequencing longer DNA strands is that this increases the longevity of the pores and yields more data out of a flowcell run. (This was the motivation of using DNA assembly for nanopore sequencing in [7]. The authors investigated the use of Gibson assembly and OE-PCR; however, the use of adaptive sampling has not been explored.)

This paper ties together the strengths and unique features of nanopore sequencing i.e., long-read PCR-free sequencing, adaptive sampling, and library recovery protocols, to provide an efficient solution to the problem of random access in DNA data storage. To demonstrate the potential of adaptive sampling, we developed a method to ligate multiple short oligonucleotides in an unordered fashion (Methods, M2). We modified the alignment algorithm in *Readfish* [8] to detect the short sub-sequences that were used for addressing different files in our experimental setup (Methods, M4). We demonstrated PCR-free random access based on adaptive sampling (Results, R1) and the ability to dynamically switch between random access targets (Results, R2), which is impossible in PCR-based random access schemes.

Further, in Results, R3, we demonstrate how using adaptive sampling for random access allows sample extraction for subsequent purification and long-term storage, so that a different file from the pool can be recovered using the same aliquot at a later time. Again, since PCR dilutes away non-requested files, this kind of information recovery is impossible with PCR-based random access schemes.

## Results

### R1. Adaptive sampling enables PCR-free random access for sufficiently long DNA oligonucleotides

To demonstrate the potential of performing random access via adaptive sampling, we prepared a test pool which consists of three “`file`(s).” The files are represented as 150-mer single-stranded DNA oligos (see Methods, M1) ligated into longer strands in random order (Figure 1) of varying length (i.e., 150, 300, 450 etc. nt, depending on the number of successful ligations) as described in Methods, M2. The resulting pool allows us to investigate the capability of ONT adaptive sampling to filter out irrelevant strands as a function of strand length. Figure 3.A (BEFORE) shows the normalised (top) and unnormalised (bottom) histograms of the number of reads observed during a sequencing run for each of the three files (labelled `file0`, `file1`, and `file2`) when `file0` is targeted. The sequencing experiments were replicated thrice; the cross-experiment variability is represented as error bars in the histograms. Among reads longer than 1500 nt (leftmost histograms), we observe that the reads almost exclusively correspond to the targeted file (`file0`), which indicates that adaptive sampling successfully performs selective retrieval of `file0`. In contrast, when no adaptive sampling is employed during a control run (Figure 3.B), the readout contains all three files, which holds across all three length cutoff levels (i.e., 1500, 750, and 500 nt, meaning that the reads shorter or equal to the corresponding cutoff level are ignored when generating the histogram).

**Fig 3.**
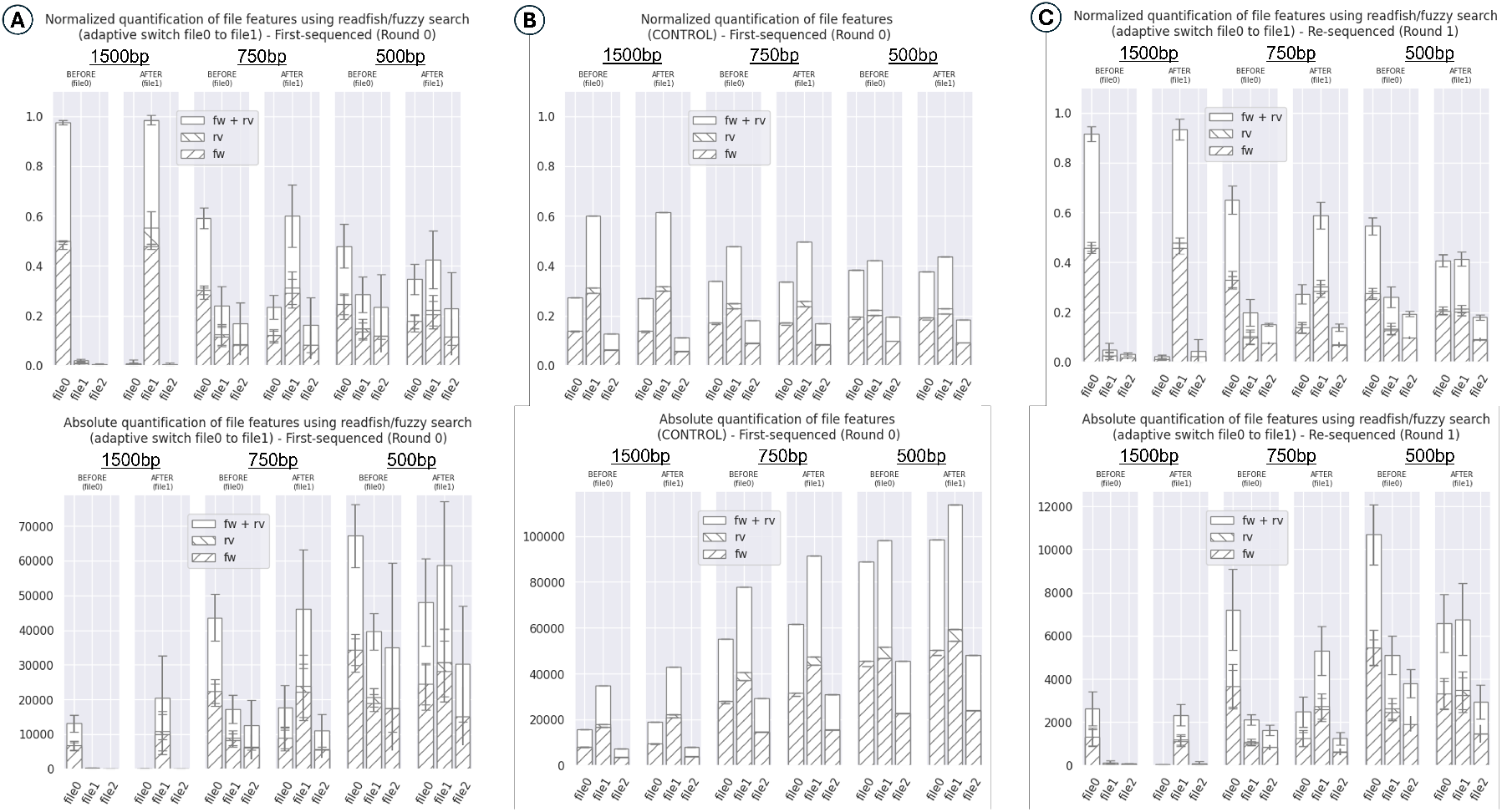
Histograms of the number of reads obtained during adaptive sampling experiments. **A)** Normalized (top) and Absolute (bottom) read count histograms for dynamic switching between `file0` and `file1` (`Round 0`). **B)** Control run. **C)** Dynamic switching after purification and re-sequencing (`Round 1`). Variance shown for `n=3` samples except the control where `n=1`.

On the other hand, adaptive sampling fails to effectively select for the targeted file (`file0`) among shorter reads. For instance, comparing the right-most (BEFORE) histograms for a cutoff at 500 nt in the adaptive sampling regime (Figure 3.A) with a control run performed without adaptive sampling (Figure 3.B), we observe no differences in relative representation of the files. This highlights that longer strands are a pre-requisite to using adaptive sampling for random access. Our ligation protocol (Methods, M2) results in progressively fewer strands of increasing length (Figure 1.C); this is reflected in the unnormalised histograms of read counts (Figure 3.A, bottom), where we observe that as the read length cutoff increases, progressively fewer reads are observed.

Overall, this experiment demonstrates that adaptive sampling can be used to effectively perform selective access to specific files in a DNA pool, provided that the pool consists of sufficiently long (*≥* 1500 nt) DNA strands.

### R2. Adaptive sampling allows dynamic switching of the targeted file

A crucial feature of adaptive sampling is that the target file can be changed dynamically during the sequencing run. To demonstrate that capability, we dynamically switched from targeting `file0` to `file1` two hours into our sequencing runs. The corresponding read histograms are reported in Figure 3.A (AFTER). We observe that the file prevalence pattern changes accordingly (especially for a cutoff length of 1500 nt). This is an interesting feature that can be used to increase the amount of useful information extracted from an aliquot of the DNA pool should additional files be requested by the user during a sequencing run. Moreover, the ability to dynamically switch adaptive sampling targets can be used to selectively target regions *within* a given file, similarly to the work in [9], to pro-actively target under-represented regions of the file and maximise the rate of information acquisition (in this setup, within-file index sequences could be used instead of file identifiers). Finally, this feature enables the use of the sample as “hot cache”: after the requested file has been recovered, the sample can be extracted from the flowcell using ONT’s library recovery protocols (see Methods, M5) and re-used for recovering other files within a short time window from the initial sequencing.

### R3. Random access via adaptive sampling enables extracting the sample for subsequent re-archiving and retrieval

After performing the sequencing experiments described in Sections R1 and R2, we extracted the sequenced sample (by liquid aspiration) from the nanopore flow cell device and purified the DNA (see Methods, M5). The extracted sample contains both sequenced (passed-through, Figure 2.A) and rejected (blocked-through, Figure 2.B) strands from the original aliquot. We then performed sample preparation again and re-sequenced the extracted sample, emulating information retrieval at a later stage after re-archiving the aliquoted pool (Figure 3.C), which is unachievable for PCR-based random access schemes (for files that were not originally targeted).

The results are shown in Figure 3.C. The figure demonstrates that adaptive sampling manages to successfully perform selective file retrieval as before; however, the non-normalised histograms (bottom) show that there are fewer oligos encountered during sequencing (about 10% of the number of reads obtained in the original experiment in Figure 3.A, bottom). The observed drop in the number of reads can be attributed to the purification method we employed (MagBeads purification, Methods M5), and it is expected that other purification methods, e.g., extraction with phenol-chloroform, would deliver a much higher output, with potential to match the initial amounts of DNA. Moreover, as it was mentioned to us by ONT in personal communication, nanopore sequencing relies on a set of motor proteins that separate the ligated double-stranded sequences into single-stranded ones as the strands pass through the pore (upon activity of the DNA helicase shown in Figure 2); hence, strand rejection (Figure 2.B) will surely proceed after the separation of the original dsDNA into two ssDNA strands facilitated by the helicase activity. If these single-stranded molecules are not actively reconstituted into double-stranded DNA, and new sequencing adapters are not re-ligated, these ssDNA strands will not be sequencing-ready in a subsequent sequencing round (see Round 0 vs Round 1 in Figure 3.A vs 3.C, respectively). This also explains why, during the ongoing sequencing run, adaptive sampling is not likely to re-capture sequences that have been already rejected, for these sequences are either single-stranded and, even in the event of reconstitution into doublestranded molecules, lack the necessary adapters. Investigating further the sequencing and re-sequencing limitations of ONT with adaptive sampling is the subject of future work to improve the possibilities for random access in DNA data storage.

## Discussion

This work demonstrates that adaptive sampling in nanopore sequencing can be used as a mechanism for random access in DNA data storage. It offers several advantages over PCR-based random access schemes, including the ability to avoid PCR bias, perform dynamic switching between random access targets, and re-archive the sequenced sample to extract, in the future, information that was not targeted originally, thus reducing pool depletion.

However, the applicability of adaptive sampling is currently limited by the minimum length of the strands—our experiments show that a length of at least 750 nt is required for adaptive sampling to effectively reject irrelevant strands. We have proposed a method to achieve the desired lengths by ligating together multiple short fragments, but this method yields progressively fewer DNA strands of increasing lengths, which leads to a loss of usable DNA product. Given the advantages of using adaptive sampling for random access, investigating reliable ways to achieve longer DNA fragments is an interesting direction of future work, especially given recent advancements in assembling gene-length products for synthetic biology research [6]. In this regard, achieving longer DNA sequences would also reduce the overhead associated with indexing the strands within a file. On the other hand, it may also be possible to reduce the minimum length of the strands for which adaptive sampling is effective by reducing the amount of buffering performed by the sequencing device prior to basecalling. Exploring this possibility is the subject of ongoing research.

Another interesting application of adaptive sampling in the context of DNA data storage is the dynamic targeting of under-represented sub-regions *within* a file to maximise the rate of information acquisition, similarly to BOSS-RUNS [9]. In other words, it could be possible to integrate adaptive sampling with error-correction decoding and dynamically identify and sample the strands that would help decoding the most. Exploring this possibility is an interesting direction of future work; however, we must acknowledge that this also requires achieving sufficiently long DNA strand lengths.

Besides the length of DNA strands, another important limitation of adaptive sampling is that it may be inefficient when targeting very small fractions of the storage pool: if the number of strands being targeted is vanishingly small compared to the overall size of the stored DNA pool, adaptive sampling will reject the vast majority of the reads when searching for the targeted file in the haystack of irrelevant information. In such scenarios, PCR-based random access is likely to be more efficient if emulsion PCR is implemented in the pipeline to avoid PCR bias [10]. Overall, it is likely that adaptive sampling is best employed in combination with other random access methods, such as fluorescence sorting [11] [12]; exploring such hybrid random access approaches is a promising direction of future research.

Overall, this work demonstrates that adaptive sampling has many advantages that cannot be matched with alternative methods. The fact that adaptive sampling can be used to eject sequences from the pores allows the operator to save a sample for later processing and terminate sequencing as soon as the quality and quantity of the sequencing output is sufficient for downstream analysis, enabling a new way in which to make DNA sequencing more sustainable, rapid and cost-effective.

Additionally, it is possible to modify the time-frames in which ONT sequencing devices, controlled using the proposed adaptive sampling scheme using *Readfish* [8], generates output files so as to significantly increase the granularity to which sequencing in real time can be realized. This opens the door to saving a significant amount of sequencing resources, maintaining the integrity of the same sample used for sequencing while enabling the use of the same sample for further downstream processing, which is particularly advantageous for research efforts during cloning procedures in the biological sciences and bioengineering, where plasmids and other genetic devices could be sequenced in real time, and the exact same sample, retrieved directly from the ongoing sequencing run, could be used for downstream cloning processes with an utmost sequence integrity. In this sense, this work highlights the significance, in the context of available technologies and methods for DNA sequencing, of being able to screen specific sequencing reads, later retrieve the same sequence used for sequencing, and further process the sequencing sample without losing it. This capability may be applied beyond DNA storage to facilitate molecular cloning pipelines in synthetic biology and bioengineering.

## Methods

### M1. Experimental oligo pool

As a starting point for our experiments, a pseudo-random DNA pool developed for [13] and available at [14] is used. The pseudo-random pool consists of 91766 sequences, each of length 150 nt, of which 110 nt are dedicated to carrying data (and are generated randomly), and 40 nt are reserved for two 20-nt primer sequences at both ends of each sequence, following the library design proposed in [1]. The ssDNA pool was synthesised by GenScript, NJ (USA), and delivered in a volume of 80 µL at 17.5 ng/µL.

The 91766 sequences were split into three files—referred to as `file0`, `file1`, and `file2`—of approximately the same size (`file0` and `file1` contain 30589 strands; `file2` contains 30588 strands). Each file uses a dedicated primer pair. The primers were used as addresses and were targeted during adaptive sampling (see Methods, M4). The 20-nt primer pairs were chosen among those developed and optimized in [1]. The 110-nt payload sequences were generated uniformly at random; then, following [1], collisions between the primers and the payloads were eliminated via the same screening procedure based on BLAST [15].

### M2. Amplification, ligation and normalization of file samples

In order to prepare the samples for adaptive sampling experiments, the following standardized procedure is proposed.

1. Double-stranding of ssDNA for ligation (see above)

2. Ligation of dsDNA files into longer DNA molecules

3. Normalization and mixture of ligated dsDNA files

4. Sample preparation for sequencing

Before the separate ligation of `file0`, `file1` and `file2` and sample mix for subsequent sequencing, double-stranding by PCR-amplification of each 150-mer single-stranded oligonucleotide stocks was used in order to enrich the DNA pool and generate a double-stranded substrate for facilitating the ligation of strands belonging to the same file. For the amplification of each file, an aliquot of 0.6 µL of a the 17.5 ng/µL stock mixture solution was amplified using Q5 Master Mix, 2X (NEB, UK) using the 5’-phosphorylated DNA primers (IDT, USA). We used the following PCR amplification protocol based on [1]: In a PCR tube, mix 10 ng of ssDNA pool (0.6 µL) with 1 µL of 100 µM of the forward primer (see primers in Table 1) and 1 µL of 100 µM of the reverse primer, 25 µL of 2*×* Q5 Master Mix (NEB, UK) enzyme mix, 20.4 µL of nuclease-free water, and 2 µL of 1.25 mg/mL acetylated bovine serum albumin. (All primers were ordered from Integrated DNA Technologies.) Vortex and spin down the resulting 50 µL mix, and place in a thermocycler with the following program: (1) 95 °C for 3 min, (2) 98 °C for 20 s, (3) 62 °C for 20 s, (4) 72 °C for 15 s, (5) go to step 2 for a total of 18 cycles, (6) 72 °C for 30 s. Purify the reaction using AMPure XP Beads at a beads-to-sample volume ratio of 1.8*×* to maximize retention of short strands.

**Table 1.**
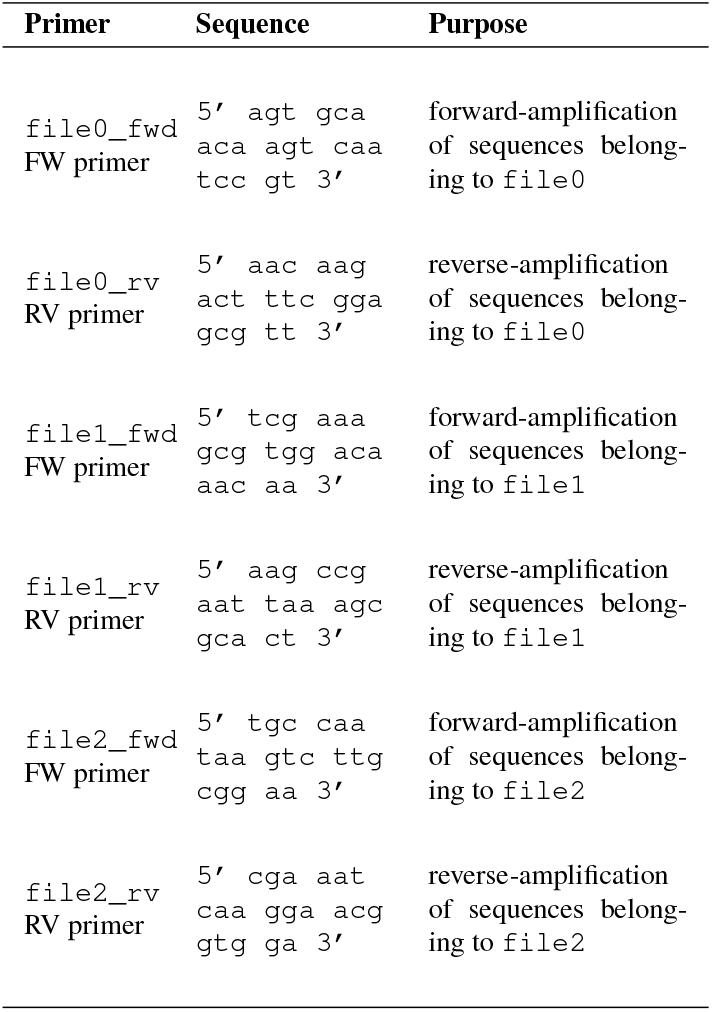
Primers used for PCR-amplification of file stocks.

| Primer | Sequence | Purpose |
| --- | --- | --- |
| file0_fwd<br>FW primer | 5' agt gca<br>aca agt caa<br>tcc gt 3' | forward-amplification<br>of sequences belong-<br>ing to file0 |
| file0_rv<br>RV primer | 5' aac aag<br>act ttc gga<br>gcg tt 3' | reverse-amplification<br>of sequences belong-<br>ing to file0 |
| file1_fwd<br>FW primer | 5' tcg aaa<br>gcg tgg aca<br>aac aa 3' | forward-amplification<br>of sequences belong-<br>ing to file1 |
| file1_rv<br>RV primer | 5' aag ccg<br>aat taa agc<br>gca ct 3' | reverse-amplification<br>of sequences belong-<br>ing to file1 |
| file2_fwd<br>FW primer | 5' tgc caa<br>taa gtc ttg<br>cgg aa 3' | forward-amplification<br>of sequences belong-<br>ing to file2 |
| file2_rv<br>RV primer | 5' cga aat<br>caa gga acg<br>gtg ga 3' | reverse-amplification<br>of sequences belong-<br>ing to file2 |

The primers used for amplification and paging of file stocks are compiled in the following Table 1.

After PCR amplification, a customized ligation protocol was performed in order to produce long, contiguous, sequences belonging to the same file. For ligation, 10 *µ*L of Quick-Ligase Buffer (NEBNext Quick Ligation Module, NEB, UK) was mixed with 0.5 *µ*L of ATP (100 mM) and 1 *µ*L of T4 PNK (NEB, UK) with a 33.5 *µ*L normalized DNA solution (approx. 2.5 ng/*µ*L) belonging to either `file0`, `file1` or `file2`, per separate. The solution was incubated at 37 °C, for 30 min, to further phosphoporylate the amplified dsDNA before the addition of 5 *µ*L of Quick-Ligase enzyme (NEBNext Quick Ligation Module, NEB, UK) and the solution was further incubated at 20 °C for 30 min.

After ligation, the resulting ligated files were separately quantified using a Qubit dsDNA HS Assay Kit (Invitrogen, CA, USA) fluorimetric assay. After quantification, the samples were mixed using a normalized ratio of 100 fmol for `file0`, 100 fmol of `file1` or and 61 fmol `file2`, respectively. The resulting mix was used for ONT sample preparation.

### M3. ONT sample preparation

All 150-mer DNA oligonucleotide preparations were cleaned-up with two consecutive steps of 80% EtOH, after mixing the sample with a 180 %(v/v) ratio of AMPure XP beads (Beckman, US). Next, a DNA dA-tailing procedure was effectuated using NEBNext® Ultra™ II End Repair/dA-Tailing Module (NEB, UK) by mixing 25 *µ*L 200 fmol of oligonucleotide sample with 3.5 *µ*L of NEBNext® Ultra™ II End Repair/dA-Tailing Buffer and 1.5 *µ*L of NEBNext® Ultra™ II End Repair/dA-Tailing Enzyme Mix before purification again with AMPure XP beads. A ligation reaction by combining 30 *µ*L of the purified end-repaired and dA-tailed oligonucleotide with 12.5 *µ*L of Ligation buffer (LNB) (Sequencing Prep Kit V14, ONT, UK), 2.5 *µ*L of Ligation Adapters (Sequencing Prep Kit V14, ONT, UK - protocol version `ACDE_9163_v114_revO_29Jun2022`) and 5 *µ*L of Quick-NEBNext Quick Ligation Module Quick Ligase (NEB, UK) incubated for 10 minutes at room temperature. After ligation, a successive modified step for AMPure XP cleanup, in which instead of the two steps of 200 *µ*L of 70 % EtOH for paramagnetic-bead clean-up, two-steps of 125 *µ*L of Short Fragment Buffer (SFB) (Sequencing Prep Kit V14, ONT, UK) is used instead, to remove excess adapters. The DNA is then quantified using a suitable method, such as a fluorometer. Finally, the prepared DNA are loaded onto the nanopore flow cell or flongle, according to the manufacturer’s instructions, ensuring proper settings on the MinION device.

### M4. Random access using adaptive sampling

We used an ONT MinION sequencing device connected to a laptop with MinKNOW and *Readfish* [8] configured following standard instructions (an interesting alternative to *Readfish* is UNCALLED [16], which directly matches raw current signals to a reference sequence and, unlike *Readfish*, avoids basecalling). By default, *Readfish* uses minimap2 [17] for mapping the reads to the user-provided reference. However, minimap2 uses alignment heuristics that are tailored to detecting alignments in long genomic reference sequences and fails to detect the presence of short 20-nt primer subsequences that we are using for addressing in the random access setup. We have therefore modified the mapping step in the implementation of *Readfish* to use the Smith-Waterman algorithm for optimal local alignment [18] [19] implemented in biopython [20] (the relatively short length of both the target sequences and the analysed read fragments allows us to perform optimal local alignment using standard dynamic programming algorithms without resorting to heuristics employed in minimap2). We used the `NUC.4.4` substitution matrix and affine gap penalties of 10 for opening and 0.5 for extending the gap. The two target primer sequences that correspond to the requested file were aligned to the read in both forward and reverse-complemented forms; a successful detection was declared if the score of the highest-scoring alignment is at least 70. The threshold score was optimised to maximise the number of sufficiently long alignments.

We remark that using the symmetric `NUC.4.4` substitution matrix implies a symmetric pattern of nanopore sequencing errors, whereas it is known that nanopore sequencing introduces transitions at a considerably higher rates than transversions [21]. Using a substitution matrix tailored to the error patterns of nanopore sequencing (or even tailoring the entire error model to the idiosyncrasies of nanopore sequencing, e.g., [13], and use that model to score alignments) should in principle improve detection accuracy. However, our analysis of adaptively sampled readouts (see Results R1) shows that the affine-gap error model with a symmetric substitution matrix performs reasonably well.

### M5. Sample extraction and purification

For the extraction and purification of DNA retrieved from the ONT flongles (R10.4.1 flow cells), used for sequencing, for a subsequence round (i.e. `Round 1` in Figure 3.C) of experimentation with sequencing and adaptive sampling, the samples were retrieved following the ONT protocol `LIR_9178_v1_revJ_11Jan2023`, mixed with a 180% (v/v) ratio of AMPure XP beads (Beckman, US), cleaned up with two consecutive steps of 80% EtOH. The resulting purified sample was prepared again using the ONT sample preparation protocol as described in Methods, M3.

### M6. Extraction of **file#** frequencies from adaptive sampling sequencing runs

For the obtaining of the `file#` count frequencies displayed in Figure 3.A, 3.B and 3.C, the *Readfish* program was used to control the ONT sequencing machinery to instruct the rejection from the pores undergoing the sequencing reads of `file` sequences not belonging to `file0` or `file1` specification, as depicted in Figure 2. BEFORE and AFTER time frames of two hours were chosen in order to clearly differentiate between the capability of the proposed scheme to discriminate between `file0` and `file1` in a sample where both `file0`, `file1` and `file2` were present. The addition of `file2` was included as a control. All sequencing runs were recorded for a period of 4 hours; adaptive sampling of `file0` was executed during the first two hours, while `file1` was screened during the last two hours of the total sequencing run.

Generated data was retrieved from from .*fast5* files that where subsequently basecalled using *dorado-0*.*8*.*2-linux-x64* ONT basecaller, to generate .*fastq* files ready for frequency analysis of the most abundant `file`(s) found at a particular time period of the sequencing run. For file frequency count, the reads listed within the .*fastq* files were extracted from the bulk files according to timestamps using the code provided in the Supplementary Section SS2.

Once the files were organized by before&qafter i.e., reads generated during the first two hours versus reads generated during the last two hours of the sequencing run, a frequency count of the number of appearances of the flanking regions encoding the priming sequences belonging to either `file0` or `file1` (which correspond to the priming sequences compiled in Table 1) were used to obtain the read counts for the histograms in Fig. 2.

## ACKNOWLEDGEMENTS

The authors are thankful to Oxford Nanopore Technologies (ONT) support staff for the continuous correspondence, assistance and troubleshooting.

## Supplementary Material

### Supplementary Section SS1: Code of the modified *Readfish* setup for adaptive sampling

The code for the modified *Readfish* setup (Methods, M4) and configuration are available via https://github.com/ImperialCollegeLondon/AdaptiveSamplingDNAseq

### Supplementary Section SS2: Code for ONT.fastq file read extraction by timestamp and read length

This code used for DNA extraction of .*fastq* file reads from ONT sequencing by timestamp and read length is also available via https://github.com/ImperialCollegeLondon/AdaptiveSamplingDNAseq/tree/main/

~~~
DNAextractor
import argparse
import glob
import os
import re
from datetime import datetime
#import pytz
def check_timestamp(line, target_time, mode):
    # Define the pattern to match the timestamp
    timestamp_pattern = r”start_time=(\d{4}-\d{2}-\d{2}T\d{2}:\d{2}:\d{2}Z)”
    print(“target_time_before”, target_time)
    target_time = str(target_time)
    target_time = datetime.fromisoformat(target_time.replace(“Z”, “+00:00”))
    print(“target_time_after”, target_time)
    # Search for the timestamp pattern in the line
    match = re.search(timestamp_pattern, line)
   # If timestamp pattern is found, extract and parse the timestamp
   if match:
      timestamp_str = match.group(1)
      #timestamp = datetime.fromisoformat(timestamp_str).replace(tzinfo=pytz.UTC)
      timestamp = datetime.fromisoformat(timestamp_str.replace(“Z”, “+00:00”))
      print(“timestamp”, timestamp, “target_time”, target_time)
      # Check the mode and perform the comparison accordingly
      if mode == “before_or_equal”:
          return timestamp <= target_time e
      lif mode == “equal_or_after”:
          return timestamp >= target_time
      elif mode == “equal”:
          return timestamp == target_time
      elif mode == “before”:
         return timestamp < target_time
      elif mode == “after”:
         return timestamp > target_time
     elif mode == “any”:
         return True
     else:
         raise ValueError(“Invalid mode. Please choose ‘before_or_equal’ or ‘equal_or_after
else:
        return False # If timestamp pattern is not found in the line
def read_dna_sequences(file_paths, target_time, mode, min_length):
    sequences = []
    for file_path in file_paths:
        with open(file_path, ‘r’) as file:
            current_sequence = ““
            in_sequence = False
            for line in file:
                line = line.strip()
                print(“Line:”, line)
                print(“Timestamp Check Result:”, check_timestamp(line, target_time, mode)) if “@v4.1.0” in line and check_timestamp(line,target_time, mode) is True:
                  in_sequence = True
                  continue
            elif “+” in line:
                 in_sequence = False
                 if current_sequence:
                  if int(len(current_sequence)) > int(min_length):
                   sequences.append(current_sequence)
                  current_sequence = ““
                 continue
             if in_sequence:
                  current_sequence += line
 print(“current sequence:”, current_sequence)
 print(“FINAL SEQ:”,sequences)
 return sequences
def write_fasta_file(sequences, output_file):
    with open(output_file, ‘w’) as file:
        for i, seq in enumerate(sequences, start=1):
            file.write(f”>Sequence_{i}\n”)
            file.write(f”{seq}\n”)
def main():
    parser = argparse.ArgumentParser(description=“Extract DNA sequences from text files in a
    parser.add_argument(“input_folder”, help=“Path to the input folder containing text files
    parser.add_argument(“output_file”, help=“Path to the output fasta file.”)
    parser.add_argument(“--mode”, choices=[“before”, “after”, “before_or_equal”, “equal_or_af
    parser.add_argument(“--target_time”, help=“Target time for comparison in the format #
    parser.add_argument(“--target_time”, required=True, help=“Target time for comparison in
    parser.add_argument(“--min_length”, default=0, help=“Target cut-off minimum length of the
    args = parser.parse_args()

   input_folder = args.input_folder
   output_file = args.output_file
   # List all text files in the input folder
   file_paths = glob.glob(os.path.join(input_folder, “*.fastq”))
   dna_sequences = read_dna_sequences(file_paths, args.target_time, args.mode, args.min_leng
   write_fasta_file(dna_sequences, output_file)
   print(“Fasta file has been created successfully!”)
if name == “ main ”:
main()
~~~

